# High-density and scalable protein arrays for single-molecule proteomic studies

**DOI:** 10.1101/2022.05.02.490328

**Authors:** Tural Aksel, Hongji Qian, Pengyu Hao, Pierre F. Indermuhle, Christina Inman, Shubhodeep Paul, Kevin Chen, Ryan Seghers, Julia K. Robinson, Mercedes De Garate, Brittany Nortman, Jiongyi Tan, Steven Hendricks, Subra Sankar, Parag Mallick

**Author notes:** To whom correspondence should be addressed Corresponding author at: 835 Industrial Rd, Suite 200, San Carlos, CA 94070, USA. Email Address (Parag Mallick).

## Abstract

Single-molecule proteomic studies are critically important for understanding the molecular origins of cellular phenotypes. However, no currently available technology can achieve both the single-molecule sensitivity and high dynamic range required to comprehensively analyze the complex mixtures of proteins in biological samples. One approach to achieve high sensitivity across a wide dynamic range would be to create a protein array that arranges billions of single molecules with regular spacing on a patterned surface. However, creating such a protein array has remained an unsolved challenge for the field. Here, we present a highly scalable method for fabricating dense single-molecule protein arrays using a specially designed DNA origami structure, protein click-conjugation, photolithography and surface functionalization. The origami-structure is enhanced via terminal deoxynucleotidyl transferase-extension, which generates brush-like projections, increasing the effective size of the origami from 88 nm to greater than 200 nm. These particles are large enough to enable super-Poisson deposition of individual protein molecules on a nano-patterned chip (>98% occupancy with only 1% of sites occupied with multiple protein molecules). This approach allowed for single-molecule protein display of 600 million protein molecules per microscope slide-sized chip with the potential to scale further with denser feature spacing. We hypothesize that this technology will ultimately enable the development of highly scalable proteomic analysis platforms that address the currently unmet need for protein measurements at single-molecule sensitivity across an exceptionally wide dynamic range of protein concentrations.

## INTRODUCTION

Tools that can quantify proteome dynamics with single-molecule resolution^1-3^ are critically needed to address a myriad of challenges in biomedicine, including the measurement of low abundance proteins in therapeutic development^4^ and biomarker discovery^5,6^ as well as for single-cell proteomics studies^7,8^. Additionally, such tools could provide insights into the molecular heterogeneity of populations of proteoforms, which is currently masked by bulk measurements. Building novel tools to measure proteome dynamics with single-molecule resolution is challenging because both single-molecule sensitivity and a high dynamic range are needed to comprehensively analyze the complex mixtures of proteins in biological samples. An ideal platform would measure tens-of-billions of individual molecules, thus enabling both single-molecule sensitivity, as well as the accurate capture of both low-abundance and high-abundance proteins. One possible approach to address this challenge is to create high-density, large-scale, single-molecule protein arrays that allow massively parallel protein detection and identification. Although development of protein arrays over the last two decades has enabled progress, especially in the context of clinical biomarker discovery^9^ current protein array technology cannot be readily scaled to allow the interrogation of billions of individual undigested protein molecules.

The current gold standard for ultrasensitive protein detection is the use of single molecule array-based digital ELISA, whereby single target molecules are captured on antibody-coated beads, enabling the detection of a few molecules in a sample^10,11^. However, these approaches rely on limiting dilution: digital measurements are possible due to a large excess number of beads over the number of target molecules in the sample, so that each bead has a small number of captured target molecules. The actual number of molecules per bead will follow a Poisson distribution and there is no way to know how many target proteins were actually captured per bead. Such techniques are therefore unsuitable for multiplexed quantification of several different proteins and are unable to achieve the dynamic range needs of complex protein sample analysis.

There are several potential ways to create a patterned surface ^12-23^. Single molecule array-based digital ELISA techniques generally use microwells created using soft lithography techniques, whereby each well can hold only one bead^10^. Photolithography techniques can create a patterned surface with uniformly separated positively charged wells where each well is occupied by a large DNA scaffold^24^. Similarly, using e-beam lithography and nanoimprinting, flat DNA origami structures can be placed on a glass surface with regular spacing^25^. However, these techniques are often challenging to execute robustly, and cost-effectively, at scale. Generally, the smaller the feature-size required, the more challenging and expensive the fabrication processes are. Consequently, to-date, no approach has combined both a high-quality, low-cost, uniformly-patterned surface with the potential to scale to billions of molecules on an array.

Since it is possible to conjugate a single protein at a specific location on a DNA origami nanoparticle structure^26^, we hypothesized that DNA origami could be used as the scaffold to create single protein arrays on patterned surfaces. A defined and consistent number of fluorescent labels can be attached to origami particles, enabling their visualization and making it possible to determine if multiple particles are co-localized. Unfortunately, many origami designs are constrained in size due to limitations in the size of the commonly used scaffolds. While these limitations can be overcome using supramolecular assemblies, fabrication of such assemblies is challenging and error prone. Here we introduce an alternative approach to generate large tile-shaped origami particles, without requiring multiple units to assemble the final structure. When coupled to an efficient process that ensures single protein conjugation to the origami structures, this approach generates a self-assembling single-molecule protein array. With further optimization, this technology will lead to proteomic analysis methods with single-molecule sensitivity and the ability to cover the extremely wide dynamic range of proteins in biological samples.

## RESULTS

### Overview of approach

We set out to design an approach whereby every protein in a biological sample is placed singly on a patterned surface (**Fig. 1**). This approach uses large DNA origami-based particles that we refer to as ‘structured nucleic acid particles’ (SNAPs) and that have a single site for protein molecule conjugation. In the overall workflow, proteins are first extracted from a biological sample using standard isolation and precipitation methods and their lysines are functionalized via an N-hydroxysuccinimide-polyethylene glycol-methyltetrazine (NHS-PEG-MtZ) crosslinker. MtZ-modified proteins are then mixed with SNAPs that carry single trans-cyclooctene (TCO) molecules. TCO and MtZ react to create a single covalent bond between the SNAP and the protein^27^. The single-stranded DNA overhangs surrounding the base-tile are extended using terminal deoxynucleotidyl transferase (TdT), which creates a peripheral brush-like structure that increases the effective size and charge of the particle^28^ (see Methods for details). We term the final resulting extended particles “enhanced” SNAPs. The negatively charged SNAPs are then deposited onto positively charged patterns on a glass or silicon surface.

**Figure 1.**
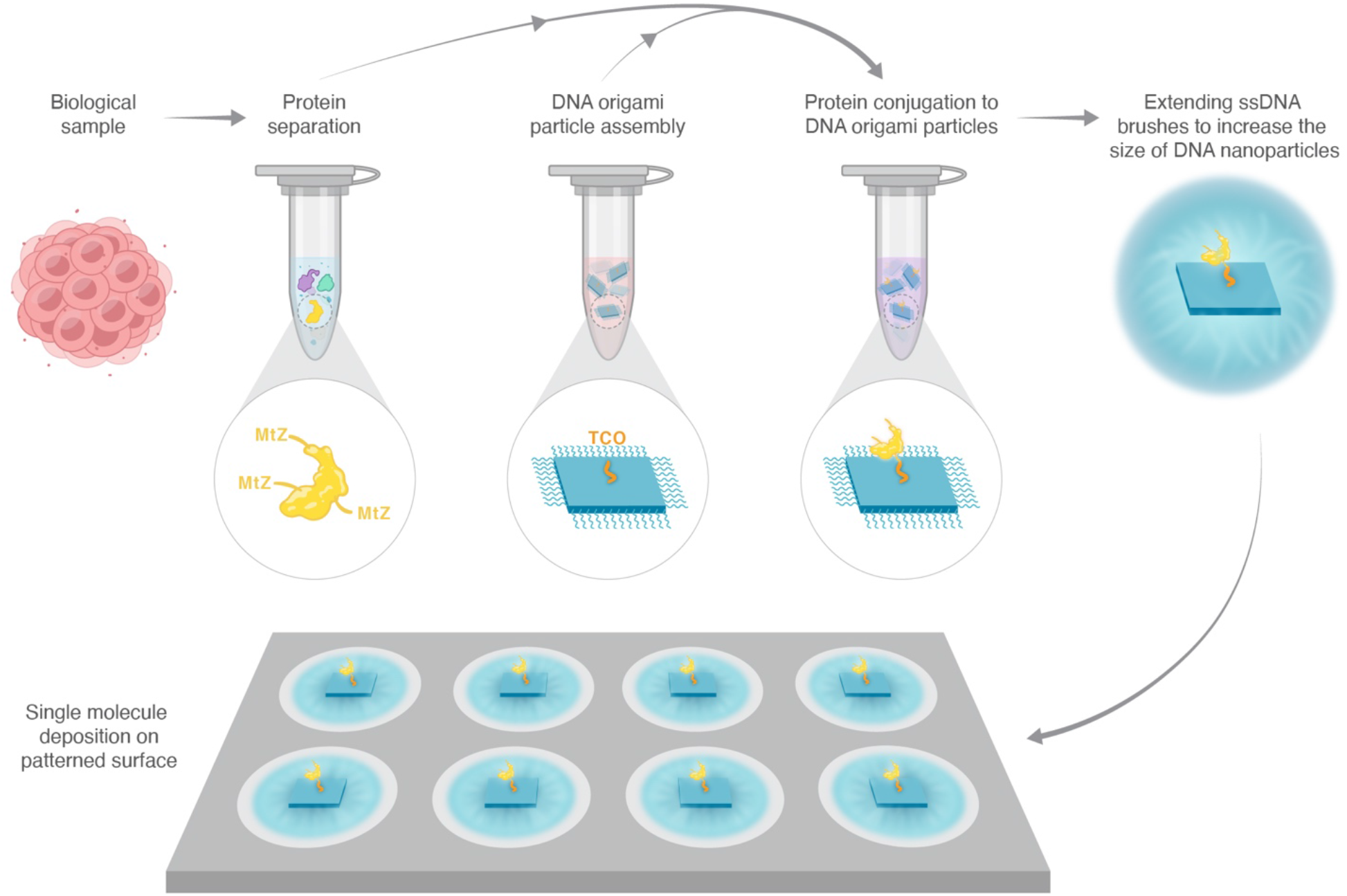
Overview of the single protein array sample prep workflow. The method presented here would be used to create single protein arrays from biological samples for single molecule studies. The workflow would consist of 1) MtZ modification of proteins separated from biological samples, 2) Production of origami tiles (SNAPs) with single TCO for click chemistry, 3) Conjugation of MtZ protein mixture to SNAPs bearing single TCO group, 4) Extension of ssDNA overhangs on the SNAP edges by terminal transferase. 5) Deposition of protein-bearing enhanced SNAPs onto APTMS functionalized patterned surface.

### SNAPs are conjugated to proteins with high efficiency

For the base component of the SNAPs, we used a single-layer, square DNA origami tile design (**Fig. 2a, SFig. 1**) with an estimated diameter of 88 nm. The DNA origami structure was folded from a mix of ssDNA oligos and the m13mp18 ssDNA. All the oligos, including the one with TCO, were mixed in excess with the scaffold DNA. Folding was achieved by heating up the mix and slowly bringing the sample to room temperature. We verified that the TCO-modified oligo in the SNAP was functional and conjugated to MtZ-modified proteins with high efficiency and specificity. Analytical HPLC results of the SNAP mixed with excess MtZ-Cy5 showed that the fraction of SNAPs containing the functional TCO group was above 95% (**SFig. 2, Table 1**). Similarly, analytical HPLC results of the SNAP, with and without the TCO group, mixed with MtZ-modified proteinA-647 showed that the protein conjugation to the TCO group was specific and the conjugation efficiency was above 95% (**Fig. 2b, Table 1**). Agarose gel analysis showed that the extension of brushes for the proteinA-647 conjugated SNAPs did not impact protein retention on the structure (**SFig. 3**). These results confirmed that TCO-modified oligos were present in more than 95% of SNAPs, and that the TCO-modified SNAPs undergo specific and efficient protein conjugation with Mtz-modified protein.

**Table 1.** TCO SNAP and MtZ reagent conjugation efficiency determined from the HPLC-SEC chromatogram absorbances.

| Conjugated molecule | Conjugation efficiency (%) |
| --- | --- |
| MtZ-Cy5 | *95±10 |
| ProteinA-647 | *96±1 |
| Ubiquitin | **90±5 |
| * Conjugation efficiencies for the fluorescent molecules to SNAP were measured as described in the methods. |  |
| ** Ubiquitin conjugation efficiency was estimated from the difference between the MtZ-Cy5 conjugation efficiency of the starting TCO SNAP and the ubiquitin-conjugated SNAP. |  |

**Figure 2.**
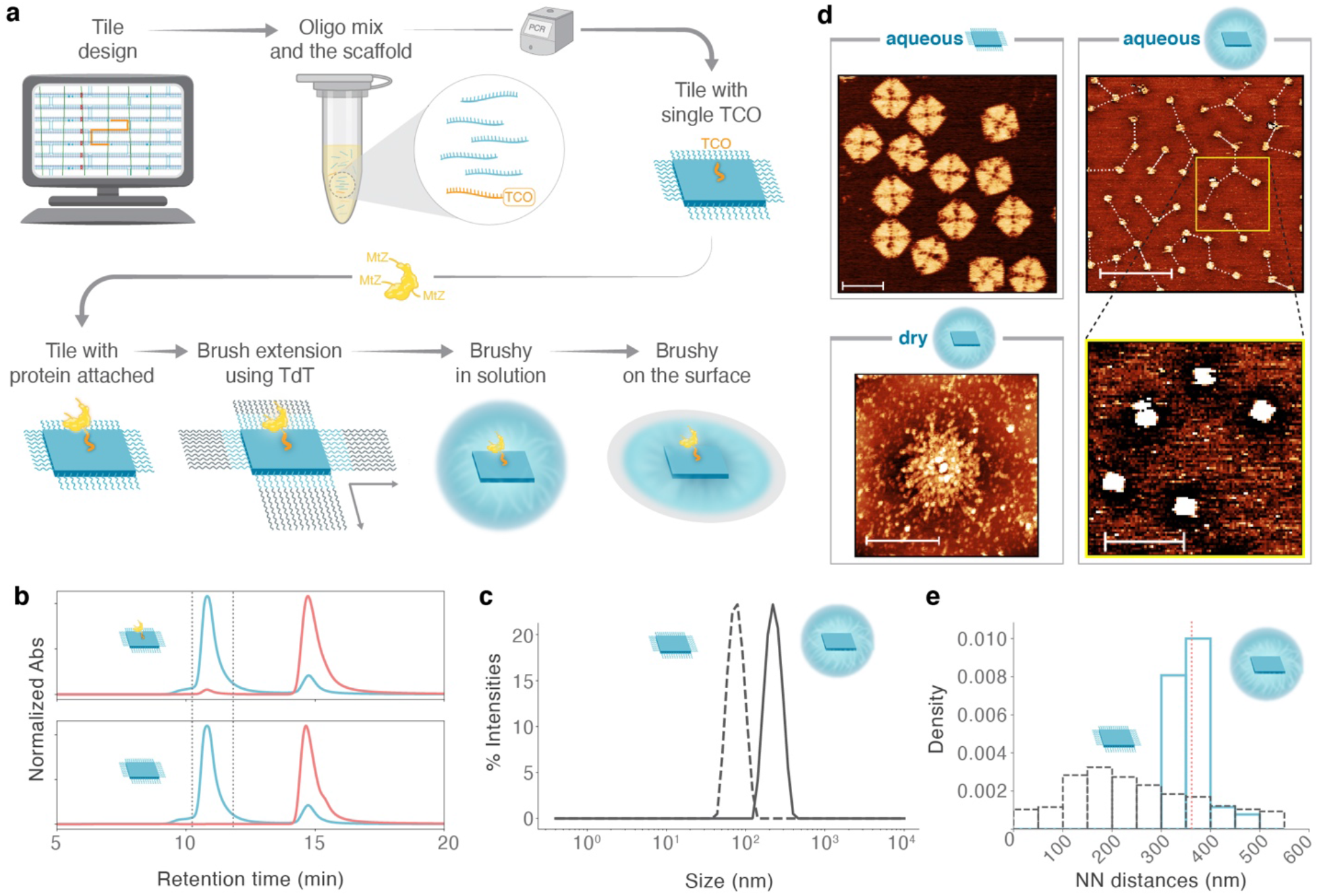
Structural and functional characterization of the SNAPs and enhanced SNAPs. **A)** Protein-conjugated enhanced SNAP production workflow. The oligo bearing the TCO moiety and the remaining staples were designed using CaDNAno (Douglas et al. 2009). The staples including the TCO-modified DNA oligo were combined with m13mp18 ssDNA scaffold in excess and the SNAP was folded over a temperature ramp on a thermocycler. The TCO-modified SNAP was coupled to MtZ protein, then the ssDNA overhangs flanking the SNAP edges were extended by terminal transferase to increase the particle size. Enhanced SNAPs were then deposited onto a patterned surface to occupy each pad with a single particle. **B)** HPLC-SEC chromatograms of TCO-bearing SNAP conjugated to proteinA-647 (top) and the SNAP with no TCO group incubated with proteinA-647 (bottom). Blue curve is for absorbance at 260 nm and the red curve is for absorbance at 652 nm. The dashed lines mark the tile peak boundaries. **C)** DLS diameter distribution for SNAP (black dashed line) and enhanced SNAP (black solid line). **D)** AFM images of SNAP and enhanced SNAP in aqueous and dry conditions. The SNAPs adopted the fold aimed at by the design (top left, scale bar is 100 nm). At saturating concentrations (3-5 nM) the enhanced SNAPs created a pattern excluding each other on the mica surface (top right, scale bar is 1 µm). The dashed lines are for the nearest-neighbor distances. The brushes were not clearly visible in aqueous imaging conditions, but we observed a halo around the enhanced SNAP particles in the phase images (bottom right, scale bar is 400 nm). When enhanced SNAPs were rinsed and dried on the mica, the full size of enhanced SNAP particles became visible (bottom left, scale bar is 400 nm). **E)** Enhanced SNAP to enhanced SNAP nearest neighbor (NN) distance distribution (blue) on mica (panel D, top right) vs. nearest neighbor distance distribution derived from simulated SNAP depositions at the same density as the enhanced SNAP density (black).

### Enhanced SNAPs are large and monodispersed

To characterize the sizes of the base-tile SNAPs and TdT-extended, enhanced SNAPs, we used both dynamic light scattering (DLS) and atomic force microscopy (AFM). DLS measurements showed that, in solution, both the base-tile SNAPs and the enhanced SNAPs were monodispersed, with an average diameter of 73.6 nm and 215 nm, respectively (**Fig. 2c, Table 2**). These measurements confirm that our “brushy” design strategy had the intended impact on the size of SNAPs. We used AFM to obtain high-resolution, single-particle structural information on the integrity and size of the particles on a 2D surface (**Fig. 2d, top-left panel**). The measured base-tile SNAP dimensions in liquid matched the expected SNAP edge length (85-90 nm) and SNAP height (1.5-2 nm) (**SFig. 4**). We imaged enhanced SNAPs on mica both in liquid (**Fig. 2d, right panels**) and as dry particles (**Fig. 2d, bottom-left panel**). In liquid, the seed tile structure was visible, but the TdT-extended brushes were not visible (**Fig. 2d, top-right panel**). In phase images, we observed a halo around the SNAPs where we expected to observe the brushes (**Fig. 2d, bottom-right panel**). When the enhanced SNAPs were dried on mica, we observed uniformly sized particles with a larger diameter of 579.8 nm **(Fig. 2d, bottom-left panel, SFig. 5)**. Given the low density of the extended arms, we did not expect to be able to accurately measure the sizes of the brushes via DLS. As such, the SNAP and the enhanced SNAP diameters measured by DLS were lower than the diameters measured from AFM images since both SNAPs and enhanced SNAPs are not perfectly spherical in 3D. Overall, these analyses confirmed that the particle structures were as expected, that brush extension did not negatively impact the SNAP fold, and that the enhanced SNAPs were substantially larger than the base-tile SNAPs, suggesting that enhanced SNAPs could exclude a larger area resulting in larger distances between the particles on a patterned surface.

**Table 2.** SNAP and enhanced SNAP diameters measured from AFM and DLS data.

| Particle type | Measurement type | Average particle diameter<br>(nm) |
| --- | --- | --- |
| Enhanced SNAP | *AFM (aqueous) | 361.0 ± 35.8 |
| Enhanced SNAP | **AFM (dry) | 579.8 ± 38.1 |
| Base-tile SNAP | AFM (aqueous) | 85.3 ± 1.8 |
| Base-tile SNAP | DLS | 73.6 ± 0.6 |
| Enhanced SNAP | DLS | 214.5 ± 6.3 |

\*Average AFM enhanced SNAP diameter in aqueous imaging was calculated from the nearest neighbor distances between the enhanced particles deposited at maximum density.
\*\* Average AFM enhanced SNAP diameter in dry imaging was estimated from the radially averaged z-height profiles. The x-axis location where the z-height flattens was picked as the effective radius of the particle.
Average AFM particle diameters were calculated from at least 10 particles. DLS diameters were averaged over 3 to 6 measurements.

To estimate the exclusion diameter for the enhanced SNAPs that created a regularly spaced pattern on mica, we measured the nearest-neighbor distances between particles. The nearest-neighbor distances yielded a tight distribution centered at 360 nm (**Fig. 2e**). To show that the observed regular pattern is unique to enhanced SNAPs, we calculated the nearest-neighbor distances for simulated SNAP depositions on mica at the enhanced SNAP density for 90 nm-wide objects. The nearest-neighbor distance distribution for simulated SNAP depositions was wider and peaked around 200 nm (**Fig. 2e**), suggesting that the observed 360 nm spacing of enhanced SNAPs on mica is a result of sub-Poissonian deposition^29^. These data suggest that the enhanced SNAPs can effectively exclude a much larger area than their central structure and suggest that they may be suitable for single SNAP deposition on chips with feature sizes of approximately 360 nm.

### Patterned chip has high single-molecule occupancy with enhanced SNAPs

To create positively-charged landing pads for high-density single-molecule deposition of enhanced SNAPs, we patterned glass wafers using photolithography and deposited 3-aminopropyltrimethoxysilane (APTMS) on the surface. We predicted that the TdT-extended brush-like structures would lay flat on the positively-charged surface. As observed from AFM images, the pad diameter and the pitch of the APTMS patterned surface averaged 370 nm with a standard deviation of 21 nm and 1.599 µm with a standard deviation of 18 nm, respectively (**Fig. 3a-c, Table 3, SFig. 6**). The patterning was composed of square regions, 200 µm by 200 µm, each containing 15,129 landing pads. A region the size of a traditional 25 mm × 75 mm microscope slide contains 37,632 of these square regions, allowing deposition of nearly 600 million protein molecules. Incubation of the patterned surface with Cy5-labeled DNA oligo yielded patterned deposition of Cy5-oligo demonstrating that the patterning and APTMS coating of the pads were successful (**Fig. 3d**). These results suggest that it is possible, using straightforward photolithography and vapor deposition techniques, to generate large-scale patterned surfaces for SNAP deposition, with tight tolerances on landing pad size and pitch.

**Figure 3.**
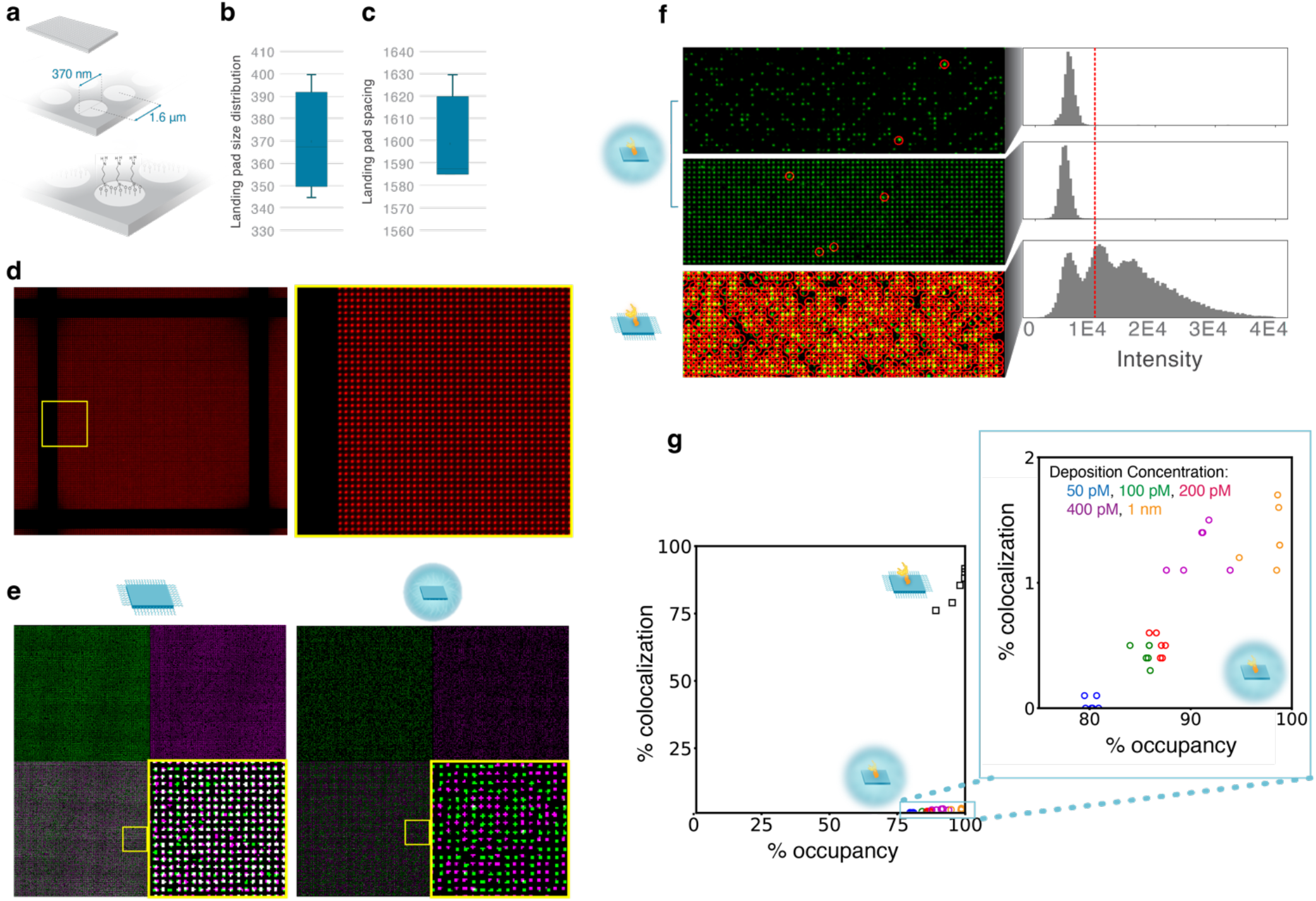
Single-molecule deposition studies with enhanced SNAPs. **A)** 2D cartoon representation for the APTMS patterned chip surface. The expected pad diameter is 370 nm and the square grid pitch on the horizontal and vertical axis is 1.6 µm. **B)** Landing pad diameter (average and standard deviation shown). **C)** Landing pad spacing (average and standard deviation shown). **D)** Alexa647-labeled DNA oligo deposition fluorescence microscopy image reveals the gridded pattern. Zoomed view of the area enclosed by yellow square is on the right. **E)** Two-color SNAP (left) and enhanced SNAP (right) co-localization images. Green is for the Alexa488 channel (top left), and purple is for the Alexa 647 channel (top right). The white pad counts on the composite image show the degree of enhanced SNAP co-localization (bottom left, bottom right is for the zoomed view). **F)** Enhanced SNAP co-localization is lower than base-tile SNAP co-localization. An increase in protein-conjugated enhanced SNAP (top: 5 pM, middle: 1 nM) deposition concentration led to higher occupancy while not causing a major impact on the intensity distributions. 50 pM deposition of the base-tile SNAP (bottom) led to a multimodal pad intensity distribution correlated with high co-localization observed with two-color enhanced SNAP mixing experiments (Panel E, left). The red circled pads on the microscopy images (left) are for the pads that have a higher intensity than the red line on the intensity histograms (right). **G)** Pad co-localization and occupancy rates. Blue is for 50 pM, green is for 100 pM, red is for 200 pM, purple is for 400 pM, orange is for 1nM enhanced SNAP deposition concentration. Base-tile SNAP (black) deposition concentration was 100 pM.

**Table 3.** Chip pad diameter and the pattern pitch data from AFM images.

| Feature | Length (nm) |
| --- | --- |
| Pad diameter | $370 \pm 21$ |
| Pitch | $1599 \pm 18$ |

Average pad diameter and pitch lengths were calculated from at least 10 measurements from AFM chip images.
Pitch is the pad-center to pad-center distance along the horizontal and vertical axis along the square grid.

To assess single-molecule landing pad occupancy on the chips, we used two-color enhanced SNAP mixing experiments (**Fig. 3e**). We produced Alexa488- and Alexa647-labeled enhanced SNAPs separately, mixed them in an equal ratio, and deposited them onto the chip at 100 pM. By counting the features that fluoresce at a single wavelength (indicating only one enhanced SNAP) and those that fluoresce at both wavelengths (indicating more than one enhanced SNAP), we assessed the degree of exclusion. Base-tile SNAP samples yielded no exclusion leading to 100% co-localization of Alexa488- and Alexa647-labeled species (**Fig. 3e left panel)**. On the other hand, at 98% deposition, enhanced SNAPs led to less than 5% co-localization suggesting that enhanced SNAPs are large enough to occupy a pad singly, excluding the deposition of a second one (**Fig. 3e right panel**) and thus leading to large-scale super-Poisson loading of the patterned surface.

Given that the experiments described above could not distinguish between single deposition and multi-deposition of the same-colored SNAPs, we additionally examined single-molecule occupancy using pad intensities. As base-tile SNAPs and enhanced SNAPs are linked with up to 40 dye molecules each, we expected the fluorescent intensities of multiple SNAPs to scale with the number of colocalized SNAPs. To identify single vs. multiple enhanced SNAPs on the pads, we performed deposition of Ubiquitin-protein-conjugated enhanced SNAPs at 5 pM and 1 nM concentrations leading to either low or high pad occupancy, respectively, and compared the intensity histograms (**Fig. 3f, top two panels**). At enhanced SNAP concentrations above 100 pM, we observed a second intensity peak with a mean and standard deviation twice that of the first peak (**SFig. 7**). At concentrations above 100 pM for base-tile SNAPs, we observed a multi-modal wide distribution indicating that there could be multiple base-tile SNAPs per pad (**Fig. 3f, bottom panel**). To quantify colocalization from single color intensities, we fitted a multimodal Gaussian distribution to the intensity histogram and quantified the contribution of multi enhanced SNAP peaks to the intensity distribution (**SFig. 7**). Using single color intensities, we verified that co-localization rates for enhanced SNAPS, even at high deposition concentrations (1 nM), were not significantly impacted by increasing deposition concentrations (**Fig. 3g**). We additionally note that protein conjugation does not have any detectable impact on SNAP deposition rate or co-localization. As seen in **Fig. 3g**, 100 pM deposition of Ubiquitin-conjugated base-tile achieved an occupancy of 98% of available features. However, the maximum average co-localization rate was approximately 90%. On the other hand, 1 nM deposition of enhanced Ubiquitin-conjugated SNAPs achieved an occupancy of 98% of available features with a maximum average co-localization rate of approximately 1.1%. Therefore, enhanced SNAPs enable high single-protein occupancy on a patterned chip.

## DISCUSSION

We have used an enhanced SNAP, fabricated using a DNA origami approach, alongside protein click-conjugation, TdT-extension, photolithography and surface functionalization to generate a scalable method for fabricating dense protein arrays with super-Poisson loading, while minimizing the number of landing pads in which multiple proteins are deposited. Additionally, it is possible to estimate which landing pad locations on the chip are likely to have multiple deposited SNAPs. Notably, the enhanced SNAPs, though not a supramolecular assembly, have diameters greater than 200 nm, and can exclude adjacent depositions such that the average distance between adjacent particles on un-patterned surfaces is approximately 360 nm. The features required to arrange such large particles are easily within reach of conventional lithographic approaches, reducing the complexity and cost of chip fabrication. We observe that this approach is not reliant upon limiting dilution to minimize multiple occupancy of a landing pad, which was approximately 1% even at 98% occupancy.

High occupancy and dense packing will ultimately enable efficient measurement by either optical or non-optical methods. We tested chips with a pitch of 1.6 µm giving approximately 600 million protein molecules per microscope slide-sized chip. Fabricating chips in a hexagonal grid with a pitch of 900 nm should be straightforward and would allow for 3x as many landing pads, thereby generating surfaces capable of holding billions of protein molecules within the space of a conventional microscope slide. Using a few such chips in concert, deposition of 10^10^ protein molecules could readily be achieved. Simulations suggest measurement of 10^10^ protein molecules would enable assays to reach dynamic ranges greater than 11 orders of magnitude, far exceeding the dynamic range of existing immuno-assays while also creating the possibility of single-molecule sensitivity.

Potential limitations of the approach will be addressed in future work. For instance, our method does not allow orientational control of the enhanced SNAPs on the APTMS functionalized landing pads via electrostatic interactions and this may kinetically impact interactions between molecules in solution and the immobilized target protein. To achieve orientational control, one side of the particles could be functionalized with a chemistry that specifically interacts or couples with the surface chemistry on the pads. In addition, the accessibility of the target protein could be further enhanced by flexible and long linkers or rigid DNA origami extensions keeping the target protein away from the SNAP surface. Another potential limitation is the lack of tight control on the size distribution of the enhanced SNAPs. The size of the enhanced SNAPs is controlled by the dTTP monomer concentration added in the terminal transferase extension reaction. Although the current process yields relatively tight distributions based on the DLS measurements and gel electrophoresis, density-based centrifugation methods could be used to fractionate the sample further to achieve a tighter distribution and potentially a lower enhanced SNAP co-localization rate.

Ultimately, we envisage that protein arrays such as those proposed above will have a wide variety of uses, including becoming a key component in the development of novel proteomic analysis methods. By immobilizing samples on the array, it may be possible to probe iteratively with a series of different affinity binding reagents to interrogate the captured proteins, in contrast to traditional bulk protein affinity measurement using a single binding event. Additionally, such arrays would enable a direct link between the identification and quantification of immobilized proteins. Further potential applications include the screening of protein-protein interactions or protein-ligand interactions, such as for compound screening. Kinetics assays could also be performed either through single-molecule imaging of interactions over time, or by using non-imaging methods such as surface plasmon resonance or impedance/conductance. Alternately, by immobilizing affinity reagents on the array, either patterned or non-patterned, it may be possible to develop ELISA-like assays with sensitivity far exceeding existing digital ELISA approaches. In summary, we expect this technology to revolutionize single-molecule proteomic studies by providing highly scalable arrays for a wide range of biological and biomedical applications.

## MATERIALS and METHODS

### Structured nucleic acid particle (SNAP) production

The SNAP was designed and modified using CaDNAno and folded as described previously ^30^. Briefly, 10 nM m13mp18 ssDNA scaffold (Bayou Biolabs, Metairie, LA) was folded with 75 nM staples (IDT, Coralville, IA). Reaction volume was 50 µl per tube. The SNAP folding buffer was 5 mM Tris-Cl, 5 mM NaCl, 1 mM EDTA, 12.5 mM MgCl2 at pH 8.0. SNAP folding was performed on Biorad Tetrad by temperature ramp starting from 90°C and ending at 20°C, decreasing one °C every minute. The folded SNAP was concentrated to nearly 100 nM using Amicon Ultra (Millipore Sigma, Burlington, MA) spin filter with 100 kDa cutoff. The SNAP was purified from excess staples using Agilent Bio SEC-5, 5 µm, 2000Å, 4.6 × 300 mm (P/N: 5190-2543) size exclusion column on an Agilent 1200 HPLC system equipped with a Diode Array Detector and Fraction Collector. Method for SNAP purification consisted of isocratic flow at 0.3 mL/min of 5 mM Tris-Cl, 200 mM NaCl, 1 mM EDTA, 12.5 mM MgCl2 (1X O.B.). Monomeric SNAP peak fractions were combined to prepare SNAPs decorated with Alexa488 or Alexa647 dye-labeled oligos (IDT Coralville, IA) and TCO-(in-house) or biotin-modified oligo (IDT Coralville, IA). Dye-labeled and TCO- or biotin-modified oligos were annealed to SNAP handles in 2X excess with respect to SNAP handle concentration. Dye oligos were hybridized to 40 complementary handles on the SNAP surface. Biotin- or TCO-modified oligo was annealed to a single complementary handle on the SNAP surface. Annealing was performed with a temperature ramp starting from 40°C and ending at 20°C, decreasing one °C every minute on Biorad Tetrad. SNAPs decorated with dye-oligos and TCO- or biotin-modified oligos were purified from excess oligos using Agilent Bio SEC-5 column on an Agilent 1200 HPLC system.

### Enhanced SNAP production

For 10 nM enhanced SNAP-3000 production, 10 nM SNAP sample was mixed with 3.6 U terminal transferase enzyme (NEB, Ipswich, MA) and 1.32 mM dTTP (NEB, Ipswich, MA) in 40 µl reaction volume. The dTTP amount required for 3000 bases extension of the 7T-3’ handles surrounding the SNAP edges was calculated via 3000 × 44 x [SNAP]. The reaction was kept at 37°C for 16 h to reach completion. The excess enzyme was removed, and the buffer was exchanged to 1X O.B. using Amicon Ultra (Millipore Sigma, Burlington, MA) spin filters with 100 kDa cutoff.

### Base-tile SNAP and enhanced SNAP-3000 characterization

SNAP and enhanced SNAP samples were run on 0.5% Agarose Gel made using Seakem LE Agarose (Lonza, Basel, Switzerland) in 0.5X TBE supplemented with 11 mM MgCl2. For visualization of the samples, SybrSafe was added to the gel from a 10,000X stock (Thermofisher, Waltham, MA). The gel was run at 70V for 3 h in an ice-water bath. The gel was visualized using Biorad Chemidoc gel imager (Hercules, CA).

Dynamic light scattering measurements of the SNAP and enhanced SNAP samples were performed on Malvern Zetasizer Nano. A 15 µl sample at around 10-20 nM concentration was added to quartz cuvette for measurements. Data was acquired with automatic measurement duration settings at room temperature.

AFM images were collected on Park systems NX-20 (Santa Clara, CA) using USC-F0.3-k0.3 (0.3 N/m) for liquid and NSC14/Cr-Au (5 N/m) for dry imaging. The data were acquired in manual mode at 1.89 Hz for high-resolution, and at 0.35 Hz for low-resolution imaging. The raw AFM images were processed using Gywddion (http://gywddion.net).

### Chip fabrication

200 mm photoresist patterned wafers were sourced from CEA-Leti (Grenoble, France). The glass (Borofloat 33, Schott) wafers were modified with an HMDS adhesion layer, followed by a lift off Layer (LOL), bottom anti reflecting coating (BARC), and photoresist stack. The photoresist was patterned in a square packed array of 320 nm round openings to reveal the silicon dioxide surface with a pitch of 1.625 microns. The patterned wafers were then modified with a water-alcohol-based plasma followed by 3-aminopropyltrimethoxysilane (APTMS) in an RPX-540 CVD system (Integrated Surface Technologies, Menlo Park, CA). A silicon monitor with 100 nm thermal oxide was placed in the chamber alongside the wafers as a proxy to measure the resulting APTMS film thickness and contact angle. Typical film thickness was about 4 Å to 5 Å and the contact angle (after an initial water rinse) ranged from 35 to 50 degrees. Following the CVD process, an additional protective layer of resist (SPR 220, Kayaku, Westborough, MA) was applied before dicing into 1-inch square die. The photoresist stack was then removed from the surface of each die by sonicating in a N-Methyl-2-pyrrolidone (NMP) bath at 50°C. The dice were then rinsed with isopropyl alcohol followed by DI water and dried under a stream of nitrogen. The contact angle was measured before and after the photoresist removal and each die was inspected under a 100X objective microscope to confirm resist removal. Representative dice were sent for AFM characterization to confirm feature size and surface roughness (Evans Analytical Group, Sunnyvale, CA). The dice were then assembled into a flow cell using a PSA gasket and a polyethylene glycol coated (Integrated Surface Technologies, Menlo Park, CA) glass slide with drilled fluidic ports (Schott).

### Flow cell assembly

The flow cell consisted of a 25 mm x 75 mm glass backer with a glass-based nanoarray chip joined by a film of pressure-sensitive adhesive (PSA). The backer had drilled holes to form the inlet and the outlet ports for the flow cell lanes. The PSA was cut in a design to define the lanes of the flow cells. The PSA layer also acted as the spacer to form the height of the channels. The flow cell assembly consisted of two main steps. First, the sub-assembly was done where the PSA was attached to the underside of the glass backer, and second, the subassembly was attached to the array chip. The PSA was placed onto the assembly fixture adhesive side facing up, using the alignment pins to register. The glass backer was then placed down on top of the PSA and pressed to complete the sub-assembly process. The assembly fixture helps align the glass backer holes to the PSA cut lanes. The final step in the flow cell assembly process was to attach the chip to the backer. The liner of the PSA was removed to expose the adhesive surface. The nanoarray chip was pressed against the backer to complete the flow cell assembly.

### TCO oligo production

5’ amine modified DNA oligo with C12 linker was purchased from IDT (Coralville, IA). Amino modified oligo was resuspended in 20% DMSO/ 0.8x PBS pH 8.0 to 2 mM. 50 molar equivalents of diisopropylethylamine (DiPEA) and 30 molar equivalents of TCO-PEG4-NHS ester (CAS 1621096-79-4) were added to the resuspended oligo in neat DMSO. Additional 20% DMSO / 0.8x PBS was added to fully solubilize all components. Final reaction composition was 1 mM amino-oligo, 30 mM TCO- PEG4-NHS ester, 50 mM DiPEA, in 59% DMSO, 0.4x PBS. Reaction was shaken at 25°C for approximately 1 h in the dark before the reaction was checked for completion (complete conversion of the amino oligo) by analytical RP-HPLC. The TCO-modified DNA oligo was purified on an Agilent 1260 HPLC system equipped with a binary pump, PDA and Fraction Collector. Purification was performed using a Phenomenex Jupiter 5 µm C5 300Å RP column (250 × 10 mm) (P/N: 00G-4052-N0). HPLC Mobile Phase A = 400 mM hexafluoroisopropanol, 16.3 mM triethylamine in water; HPLC Mobile Phase B = 100% methanol. Column temperature = 30°C. Fractions were collected in 16 s intervals from 20 to 30 min. HPLC method was as follows:

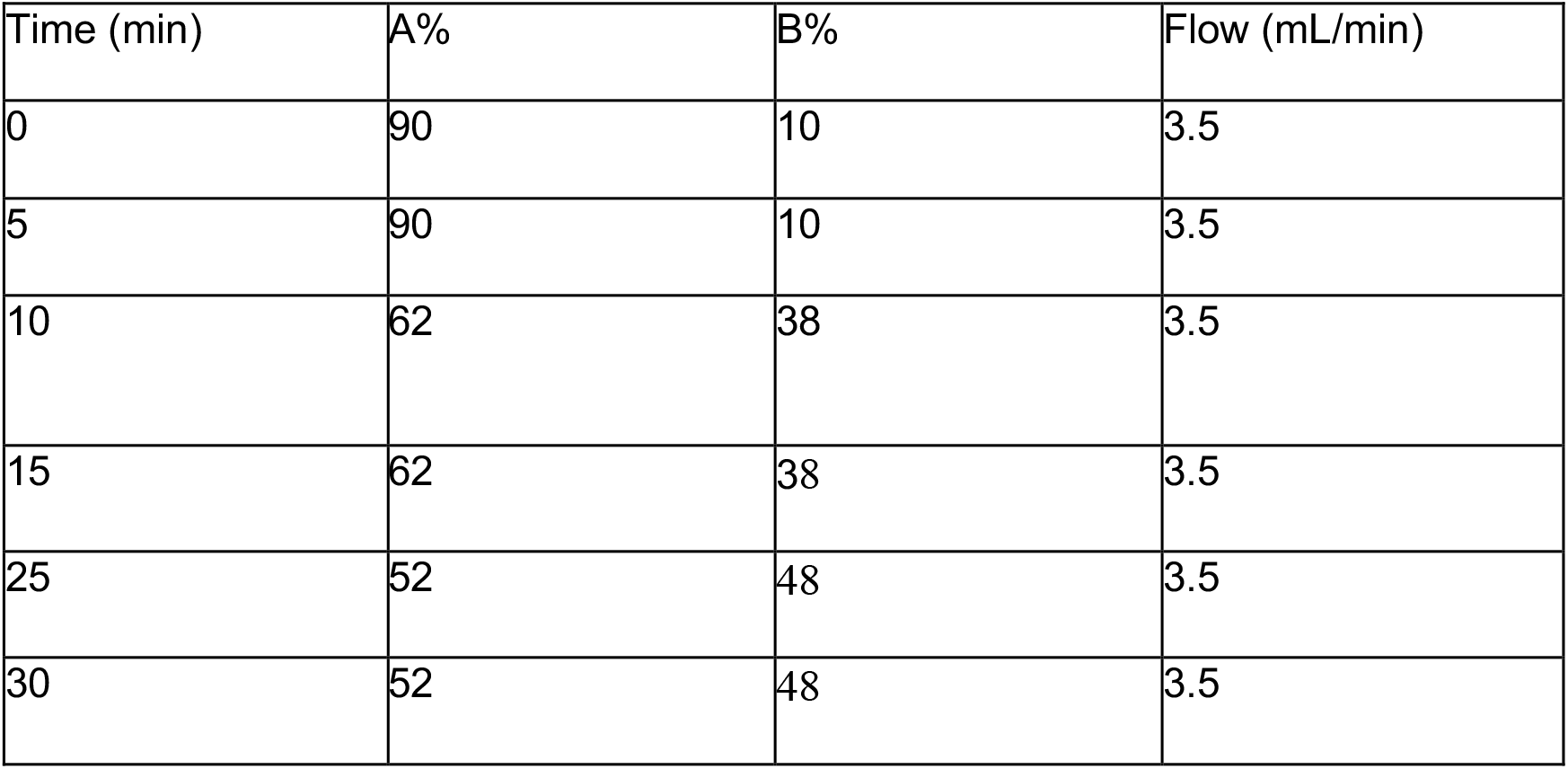
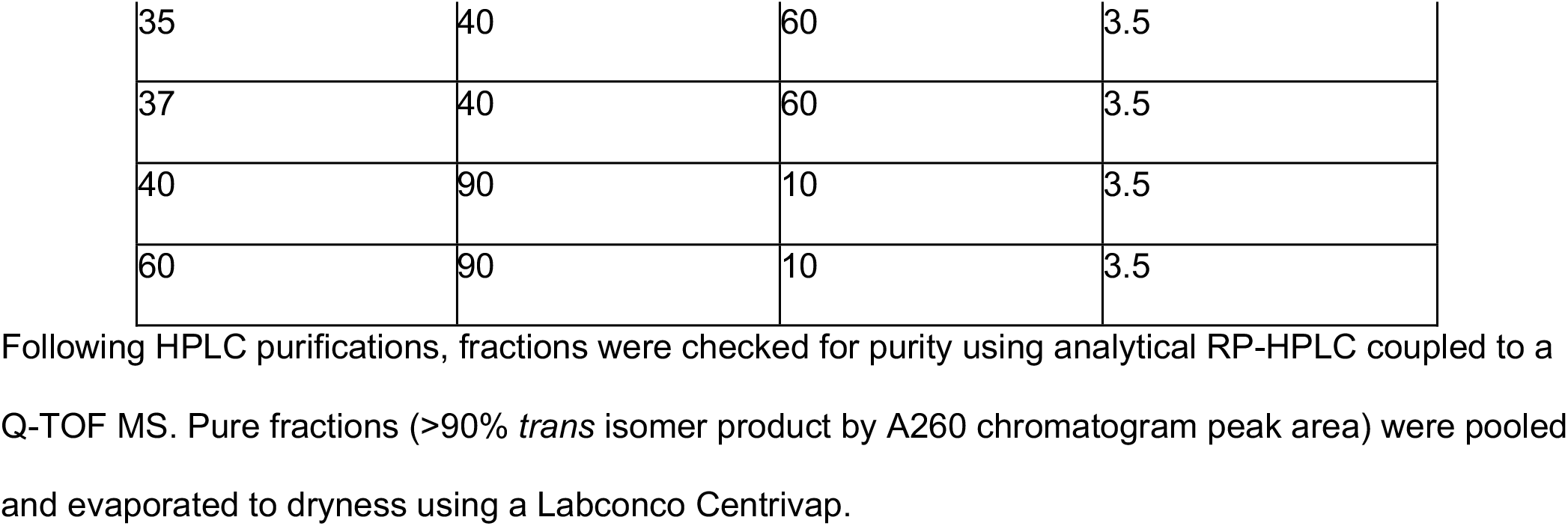

| Time (min) | A% | B% | Flow (mL/min) |
| --- | --- | --- | --- |
| 0 | 90 | 10 | 3.5 |
| 5 | 90 | 10 | 3.5 |
| 10 | 62 | 38 | 3.5 |
| 15 | 62 | 38 | 3.5 |
| 25 | 52 | 48 | 3.5 |
| 30 | 52 | 48 | 3.5 |
| 35 | 40 | 60 | 3.5 |
| 37 | 40 | 60 | 3.5 |
| 40 | 90 | 10 | 3.5 |
| 60 | 90 | 10 | 3.5 |

### Modification of ubiquitin and proteinA-647

Recombinant ubiquitin with N-terminal Histag (BPS Biosciences, P/N: 79293) and Alexa647-proteinA conjugate (Thermofisher, Waltham, MA) were modified with methyltetrazine-PEG4-NHS ester (mTz) using the following conditions. Ubiquitin was buffer-exchanged to 1x PBS pH 7.4 using BioRad Econo-Pac 10DG Desalting Columns (Biorad, Hercules, CA); 10 molar equivalents of mTz were added (from a stock 10 mM solution in dimethylformamide (DMF)), along with additional 1x PBS pH 7.4, to the protein such that the final protein concentration was 40 µM in a final solvent composition of 4% DMF in PBS pH 7.4. Reaction was incubated for 3 h at 25°C in the dark. Modification reaction was quenched by adding a sufficient volume of stock 800 mM L-Arginine in 1x PBS (not pH adjusted after arginine addition) such that the final concentration of arginine in the reaction mixture was 50 mM. Quench was performed for 2 h at 25°C, in the dark. Alexa647 protein A conjugate was modified and quenched with mTz in a similar fashion but with a final protein concentration (in the modification) of 80 µM and 30 molar equivalents of the mTz-PEG4-NHS ester, 8% DMF in 0.92x PBS pH 8.0 final solvent composition. Quenched modification reactions were purified using the manufacturer’s procedure for desalting on BioRad Econo-Pac 10DG Desalting Columns (Biorad, Hercules, CA) to remove excess mTz modifier. 10DG columns were equilibrated and eluted with 1x PBS, 200 mM L-Arginine pH 8.7. Purified proteins were concentrated to >15 µM using Sartorius Vivaspin 3k (ubiquitin) or 10k (proteinA) 3 mL centrifugal spin filters with hydrosart membranes. The concentrated proteins were rinsed in the spin filters with an additional 2 mL of 1x PBS, 200 mM L-Arginine pH 8.7. Modified proteins were stored at 4°C or directly taken to the conjugation step.

### Conjugation of MtZ-modified proteins to SNAPs

SNAPs with a single TCO (*trans-*cyclooctene) functional group, at a minimum of 100 nM, were incubated with approximately 3 molar equivalents of mTz-modified proteins in 1x O.B. at 25°C overnight (approximately 16 h) on a tube-shaker protected from light with foil. Protein-conjugated SNAP sample was purified from excess unconjugated protein via HPLC-SEC (Size Exclusion Chromatography) using an Agilent 1200 HPLC system equipped with DAD and fraction collector. The SEC column used was an Agilent Bio SEC-5, 5 µm, 2000Å, 4.6 × 300 mm (P/N: 5190-2543). Method for purification consisted of isocratic flow at 0.3 mL/min of 1x HPLC O.B. (200 mM NaCl, 11 mM MgCl2, 5 mM Tris-HCl, 1 mM EDTA) for 1.5 column volumes. Fractions corresponding to the SNAP peak were pooled and concentrated using Amicon Ultra 100k MWCO Regenerated Cellulose Spin Filters. Purified, concentrated sample was stored in DNA Low-Bind tubes at 4°C.

### Quantification of TCO oligo and MtZ reagent conjugation efficiency

Multiple tools were used to quantify the conjugation efficiency of MtZ reagents to SNAPs with a single TCO group.

For fluorescent MtZ reagents, HPLC-SEC chromatography was used to derive the relative absorbances of SNAP at 260 nm and the conjugated fluorescent molecule’s absorbance at its max absorbance wavelength. The 260 nm SNAP extinction coefficient used to quantify the relative molar ratio of SNAP to the conjugated particle was 116,898,548/ (cm*M). For Mtz-Cy5 conjugation, the extinction coefficient at 652 nm was 250,000/ (cm*M). For Mtz-ProteinA-647 conjugation, the extinction coefficient at 652 nm was 956,000/ (cm*M). The conjugation efficiency was derived using Equation 1.

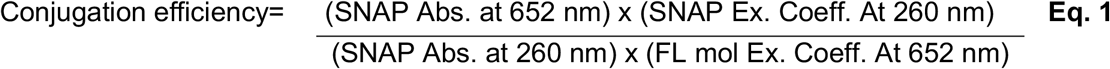

For non-fluorescent protein targets, the conjugation efficiency was assessed by the difference in MtZ-Cy5 conjugation efficiency of the starting SNAP with TCO group and SNAP coupled to MtZ-modified protein. In addition, a protein conjugated SNAP sample was run on denaturing SDS-page gel to observe the protein conjugated oligo band.

### SNAP and enhanced SNAP fluorescent imaging

In each lane of a dry patterned chip, SNAPs and enhanced SNAPs with or without the conjugated protein were deposited at 50-400 pM in running buffer (10 mM HEPES, 120 mM NaCl, 10 mM MgCl2, and 5 mM KCl, pH 7.4, 0.1% Tween-20 (Sigma P/N 44112) and 0.001% Lipidure (NOF P/N: CM5206)) for 15 min at room temperature. Excess nanoparticles were washed with 1mL of running buffer. In lanes where dark SNAPs or enhanced SNAPs were deposited, 1 µM of dye-labeled oligos (18.1 in sequence, Alexa Fluor 488 or Alexa Fluor 647) in hybridization buffer (running buffer + 100mg/mL dextran sulfate) were incubated for 2 min before rinsing out by 1 mL of running buffer. Images were taken on a home-built fluorescence microscope. The microscope objective was Olympus UCPLFLN 20x, 0.7 NA, with a correction collar for thick cover glass. The camera was Photometrics Prime BSI (2048 × 2048 pixels, 6.5 mm pixel size, sCMOS). Effective pixel size in the focal plane of the microscope objective was 0.325 mm. The excitation light source was a Necsel NovaLum 3-laser module (488 nm, 525 nm, 638 nm). The three lasers were independently controlled. The fluorescence filter set was Chroma 89402 (DAPI/FITC/TRITC/Cy5).* Laser light was focused to a 2048-pixel x 200-pixel (666 mm x 65 mm) rectangular area that was galvo-scanned (galvo scanner from Thorlabs GVS201) across the camera’s field of view synchronously with the camera’s rolling shutter. A B&K Precision 4052 waveform generator generated the sawtooth wave that controlled the galvo. The camera’s exposure time was set so that 200 rows of the sCMOS sensor detected light simultaneously. These 200 rows were in effect a confocal slit that moves across the 2048 rows of the sCMOS sensor at a rate of 1 row per 11.28 ms when the camera is operated at its maximum line rate. The line rate was reduced to 1 row per 112.8 ms when increased sensitivity was required. A total of 30 images in a 2-by-15 layout was acquired for each lane of the flowcell.

### Image processing

The input images were corrected for spherical aberration using the patterned surface array dimensions and the pad pitch as ground truth. Patterned array and single pad boundaries were determined using the subarray fluorescent patterns excited by the 488 laser. The mapping between 488 and 635 laser acquired images was constructed using a SNAP decorated with 44 Alexa488 and 20 Alexa647 modified oligos. The occupancy of the pads was determined based on the difference between the pad intensities and the background intensity. Briefly, the whole image was normalized against the background, where the background intensity estimate of the images was calculated by tiling the image (64×64 pixel) and computing the 5^th^ percentile for each tile. The computed background intensity was fitted with paraboloid. The image was then normalized against the fitted background. The primary feature score was computed for each candidate object by convolving the pads with a 5×5 gaussian kernel on the normalized image. Occupancy for each pad was classified using a one-feature clustering method to determine threshold the occupancy status of the candidate pad.

### Quantification of enhanced SNAP co-localization from fluorescent microscopy images

Two types of experiments were used to estimate enhanced SNAP co-localization on the pads. For the first experiment, Alexa488 and Alexa647 labelled enhanced SNAPs were mixed at equimolar ratio and deposited on the chips. The co-localization rate was estimated by the ratio of pads occupied by Alexa488 and Alexa647 enhanced SNAPs and all occupied pads. For the co-localization analysis, six microscopy images taken at the center of the flow cell away from fluidic inlet and outlet were used. For the SNAP co-localization analysis, we used the double color mixing data, because the SNAP single color deposition intensity distribution is wide multi-modal, not allowing easy separation of single and multi-tile peaks. For the second experiment, Alexa488 labelled enhanced SNAPs were deposited onto the surface and the co-localization rate was estimated from the fluorescence intensity histogram analysis. In this analysis, the bimodal gaussian distribution was fitted to the intensity distribution where the mean intensity and the standard deviation of the second peak were constrained to twice the mean intensity and the standard deviation of the first peak, respectively. In most of the intensity distributions with enhanced SNAPs, the second peak in the distribution had low amplitudes; therefore, constraining the mean intensity and the standard deviation of the second peak provided robust results that are comparable among images from the same experimental condition and images across different experiment conditions. The co-localization rate was quantified by Equation 2.

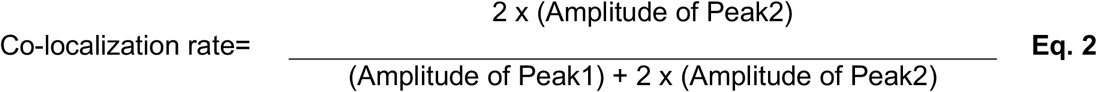

## 1. Supplementary Figures

**SFigure 1.**
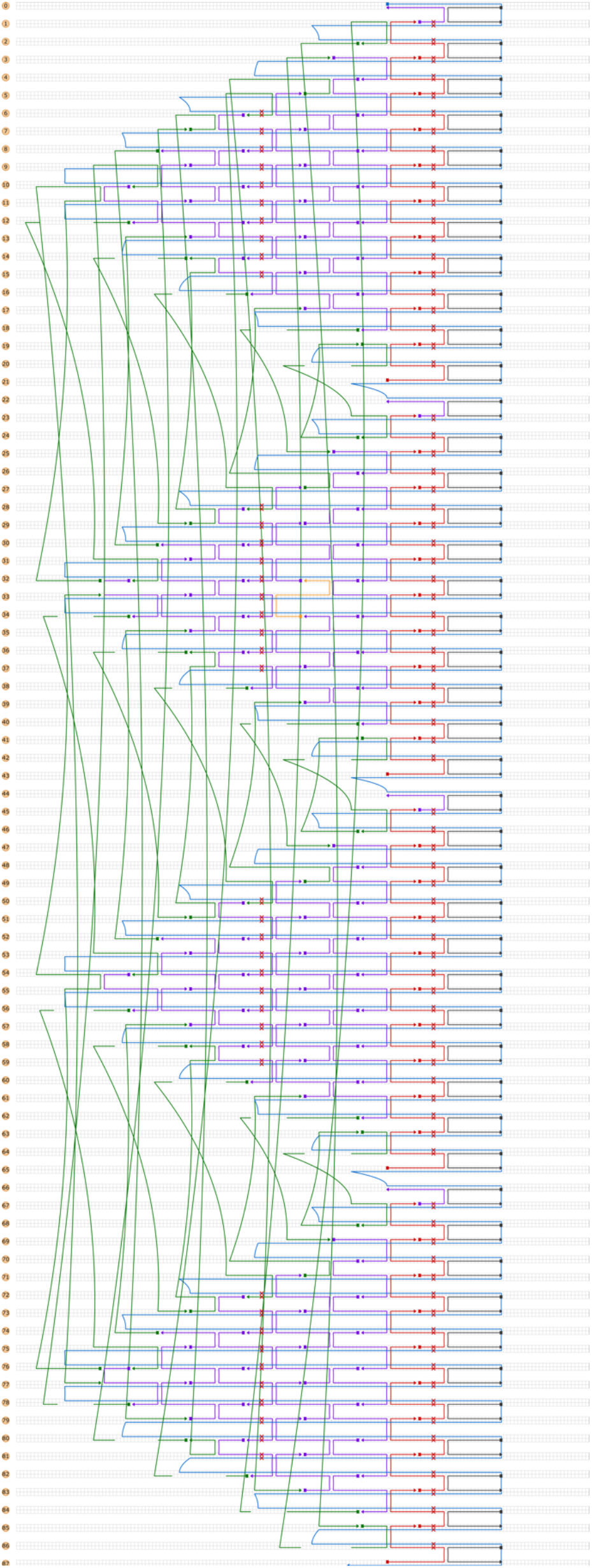
CaDNAno design diagram for DNA origami base-tile.

**SFigure 2.**
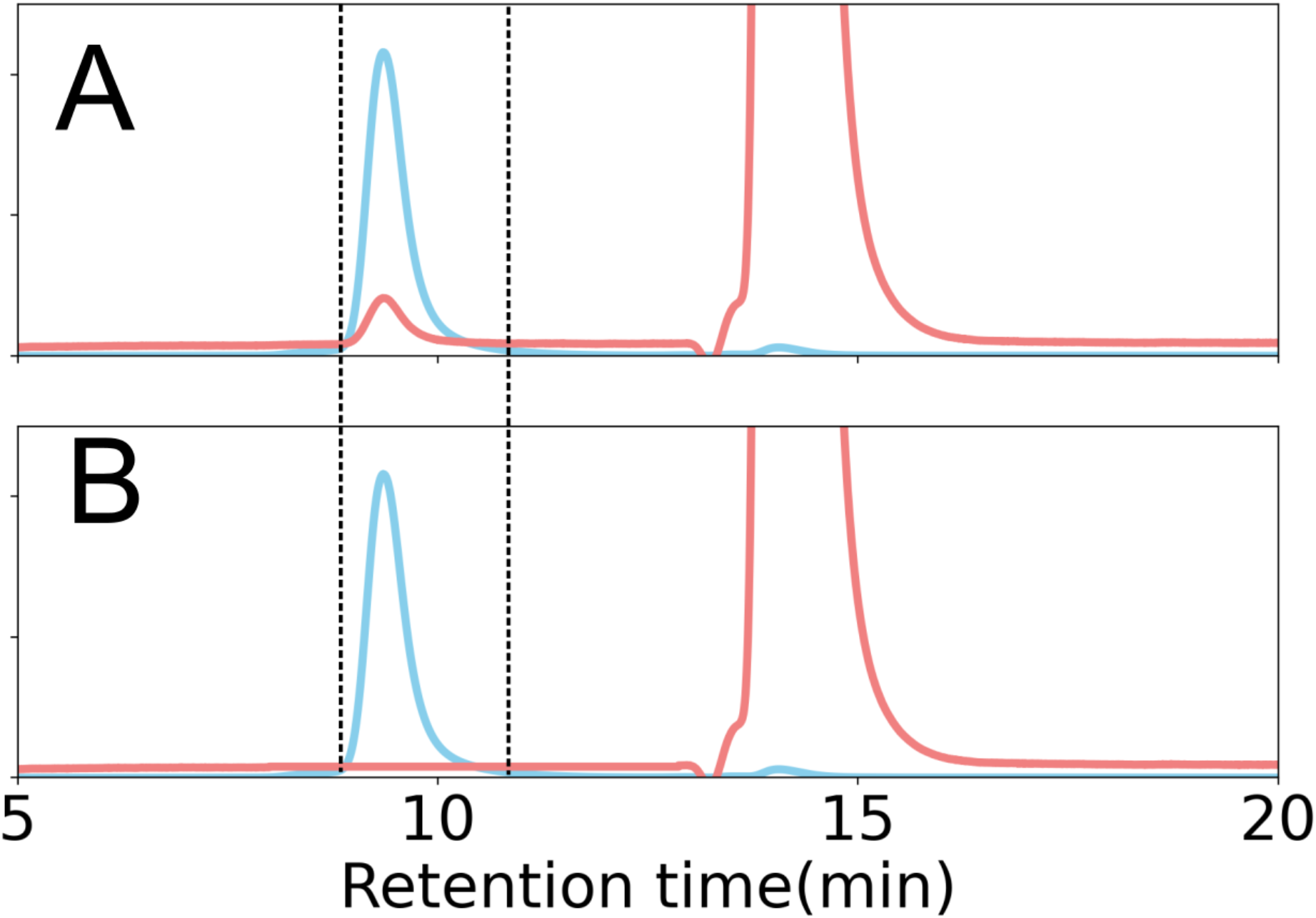
HPLC-SEC chromatograms of the (A) TCO-modified SNAP and (B) ubiquitin-conjugated SNAP incubated with Mtz-Cy5. **A)** TCO-modified SNAP incubated with excess MtZ-Cy5. The peak between the dashed line is for the SNAP. Blue is for absorbance at 260 nm and red is for absorbance at 652 nm. The right peak is for the free MtZ-Cy5. The conjugation efficiency of the TCO with MtZ-Cy5 is quantified from the ratios of the 260 nm and 652 nm absorbance peak areas. **B)** Ubiquitin-conjugated SNAP incubated with excess MtZ-Cy5. Ubiquitin-conjugated SNAP does not absorb at 652 nm suggesting that there is no SNAP with free (unconjugated) TCO, therefore all the SNAPs are conjugated to ubiquitin.

**SFigure 3.**
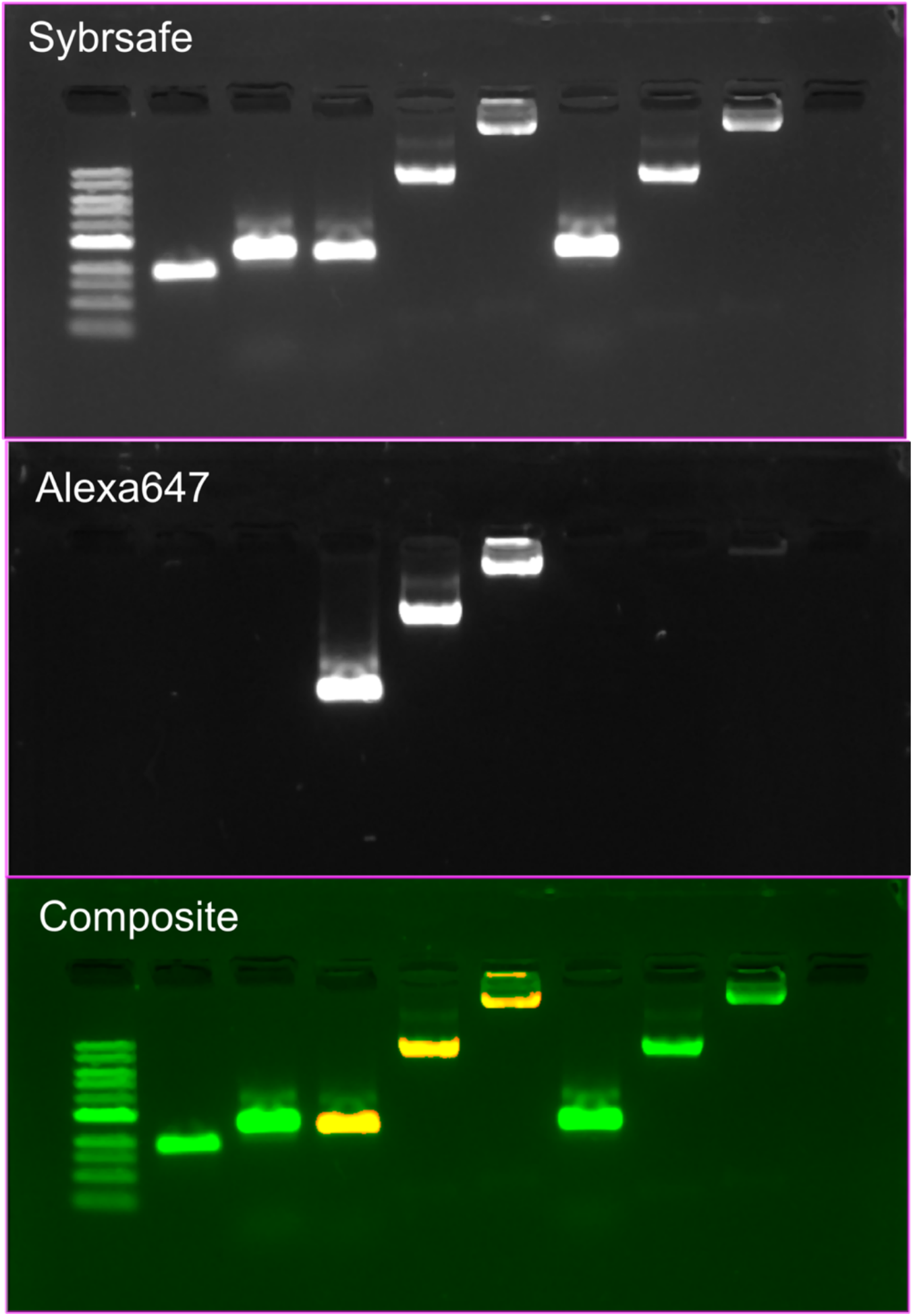
0.5% Agarose gel image for proteinA-647-conjugated SNAP and enhanced SNAP. Samples from left to right are 1) 1 kb NEB DNA Ladder, 2) m13mp18 scaffold, 3) TCO-SNAP, 4) TCO-SNAP conjugated with MtZ-proteinA-647, 5) Sample 4 extended with 1000 fold excess dTTP per initiation site, 6) Sample 4 extended with 2000 fold excess dTTP per initiation site, 7) Sample 4 extended with 3000 fold excess dTTP per initiation site, 8) Control SNAP incubated with MtZ-proteinA-647 and purified by HPLC-SEC, 9) Sample 8 extended with 1000 fold excess dTTP per initiation site, 10) Sample 8 extended with 2000 fold excess dTTP per initiation site, 11) Sample 8 extended with 3000 fold excess dTTP per initiation site. Top image is acquired with SybrSafe laser and filter settings. Middle image is acquired with Alexa647 laser and filter settings. Bottom image is the composite of the top (green) and middle (red) images.

**SFigure 4.**
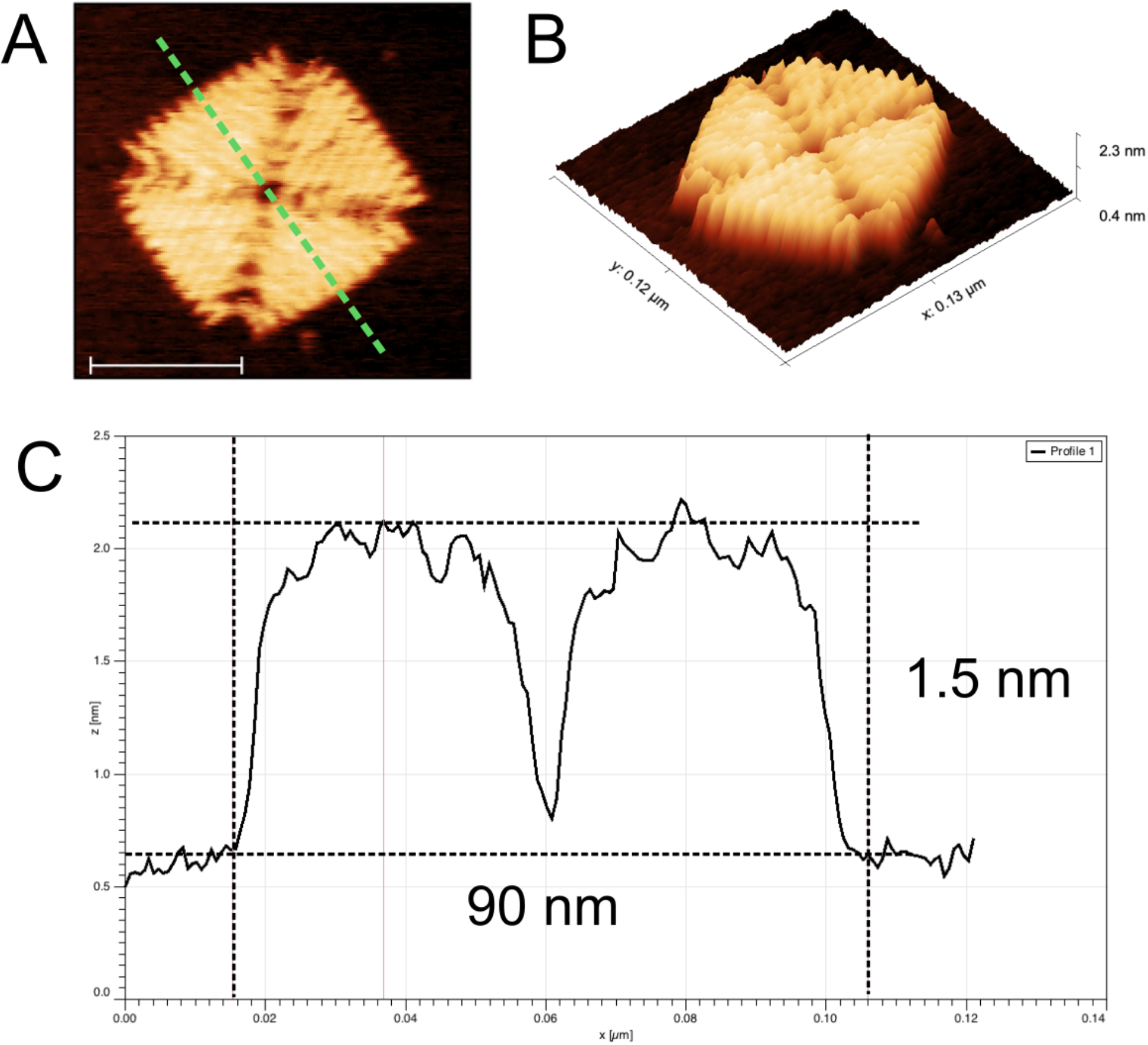
SNAP dimensions from AFM in aqueous conditions. **A)** High-resolution zoomed-in view of a SNAP by AFM. The scale bar is 50 nm. The z-height profile of the SNAP along the dashed line is in C. **B)** 3D view of the SNAP in A. **C)** The z-height profile of the SNAP in A. The length of the SNAP along the dashed line is 90 nm. The maximum height is 1.5 nm.

**SFigure 5.**
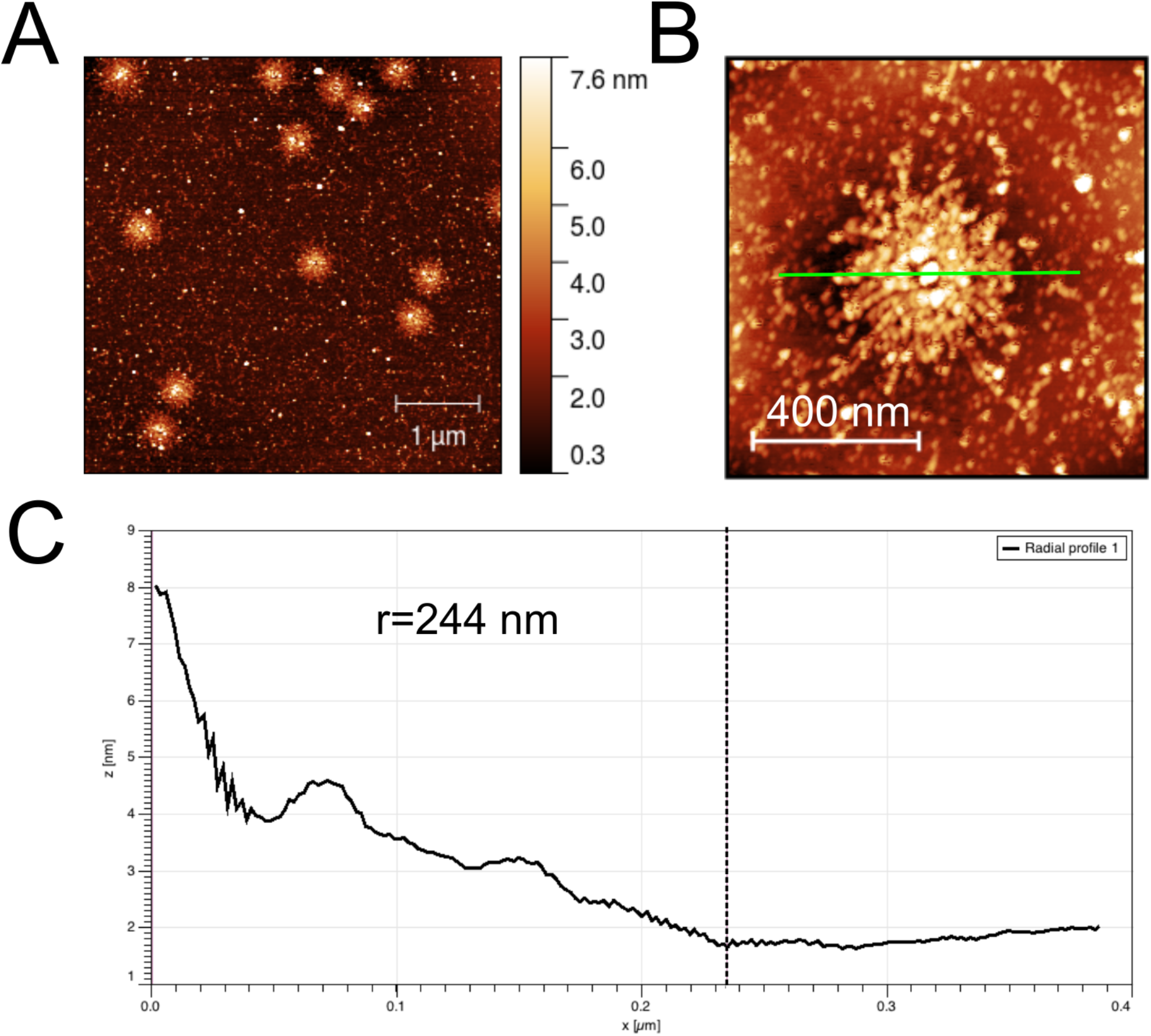
Enhanced SNAP dimensions from AFM in dry condition. **A)** AFM image of enhanced SNAPs in dry condition. **B)** Zoomed-in view of an enhanced SNAP. The radially averaged profile of the enhanced SNAP along the green line is in C. **C)** Radially averaged z-height profile of the enhanced SNAP. The radius at which the z-height flattens is marked by a dashed line. The effective diameter of the enhanced SNAP structure in B is estimated by multiplying the radius (244 nm) by two.

**SFigure 6.**
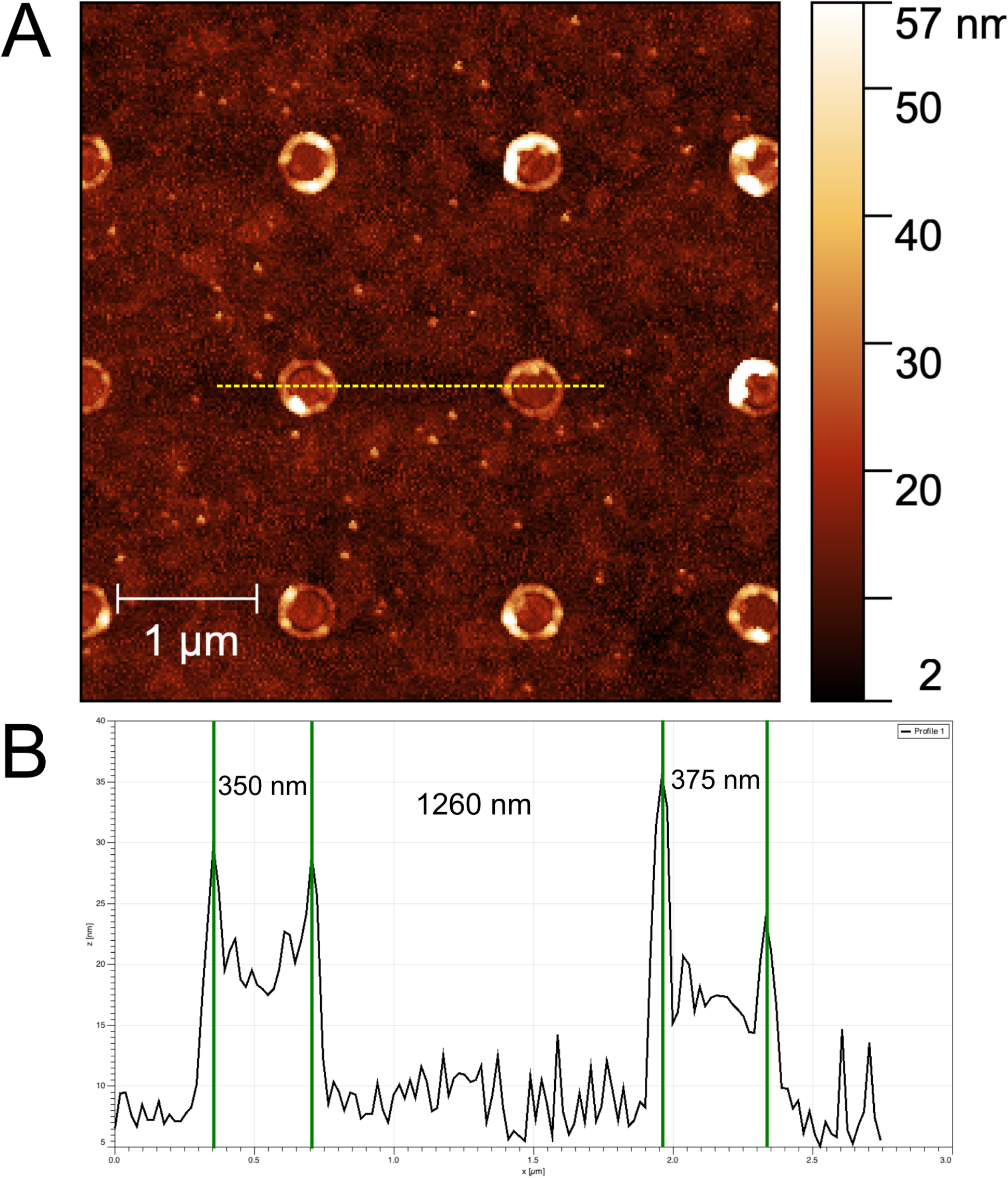
Patterned chip pad size and array pitch characterization. **A)** AFM image of the patterned chip surface. APTMS functionalized pads are enclosed by circular rims. **B)** Height profile along the green dashed line in A. The numbers show the width of the pads and the inter-pad distance in nm along the line.

**SFigure 7.**
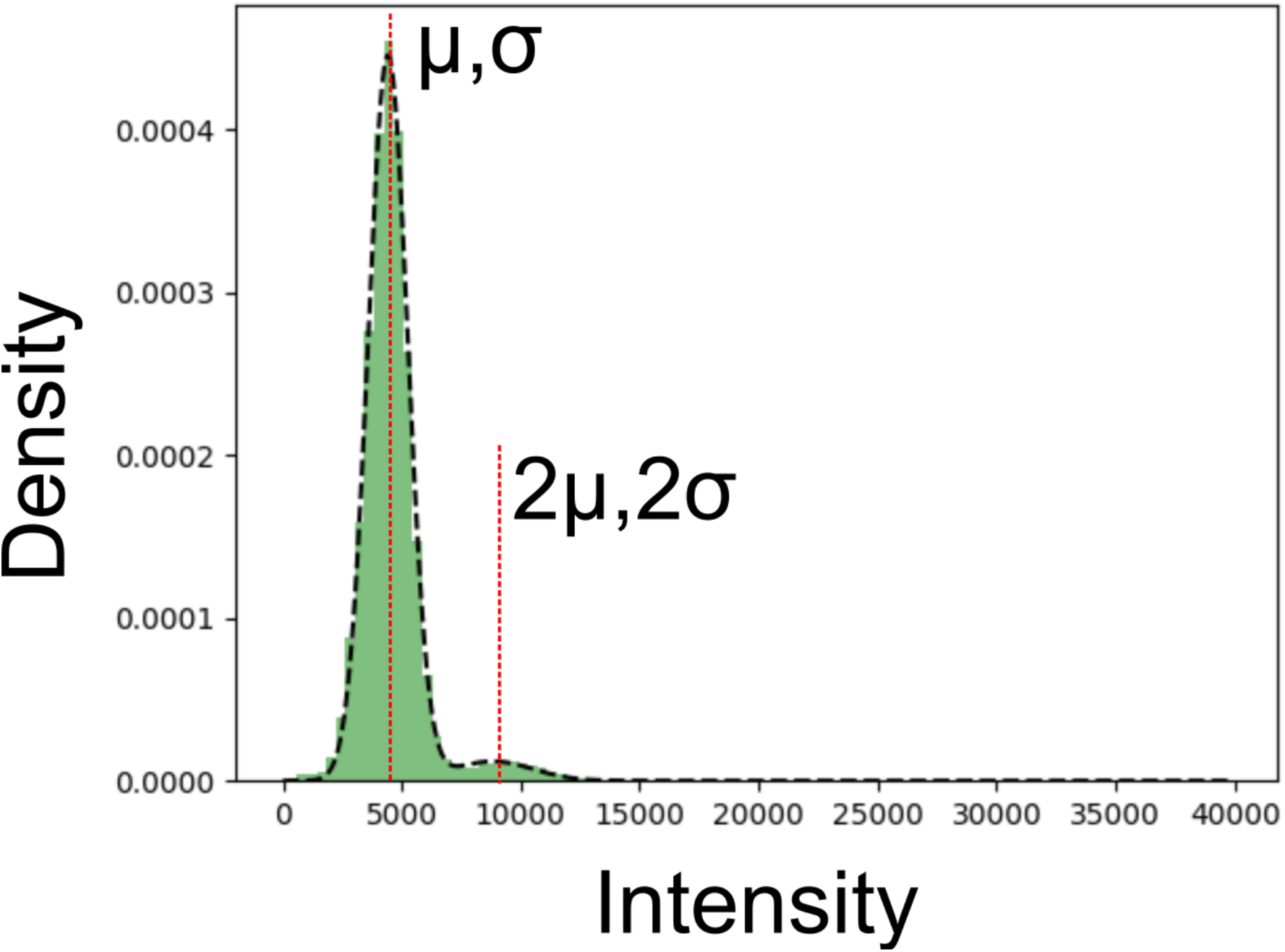
Quantification of enhanced SNAP co-localization per pad on the patterned chip from fluorescence intensities. At concentrations higher than 100 pM, the intensity distribution becomes multimodal with two distinct peaks. The enhanced SNAP titration results suggest that the second minor peak with higher mean intensity corresponds to pads occupied with two enhanced SNAPs. The co-localization rate is quantified from two modal gaussian fit to the intensity distribution where the second gaussian is constrained to have twice the mean intensity and the standard deviation of the major peak.

